# Seneca Valley virus intercellular transmission mediated by exosomes

**DOI:** 10.1101/2020.02.14.950345

**Authors:** Keshan Zhang, Guowei Xu, Shouxing Xu, Xijuan Shi, Chaochao Shen, Junhong hao, Minhao yan, dajun zhang, Xiangtao Liu, Haixue Zheng

## Abstract

Exosomes are cup-shaped vesicles that are secreted by cells and are involved in the intercellular transport of a variety of substances, including proteins, RNA, and liposomes. Studies have shown that pathogenic microorganisms are contained in exosomes extracted from pathogenic micro-infected cells. The Seneca Valley virus (SVV) is a non-encapsulated single-stranded positive-strand RNA virus that causes ulceration in the pig’s nose, the appearance of blisters, and other clinical symptoms similar to foot-and-mouth disease (FMD). Whether exosomes from SVV-infected cells can mediate SVV intercellular transmission is of great significance. There have been no studies showing whether exosomes can carry SVV in susceptible and non-susceptible cells. Here, we first extracted and identified exosomes from SVV-infected IBRS-2 cells. It was confirmed that replication of SVV can be inhibited when IBRS-2 cells treated with exosomes inbihitor GW4869. Furthermore, laser confocal microscopy and qRT-PCR experiments were performed to investigate whether exosomes can carry SVV and enable the virus to proliferate in susceptible and non-susceptible cells. Finally, exosome-mediated intercellular transmission can not be completely blocked by SVV-specific neutralizing antibodies. Taken together, this study showed that exosomes extracted from the SVV-infected IBRS-2 cells can carry SVV and transmit productive SVV infection between SVV susceptible and non-susceptible cells, this transmit infection is resistant to SVV specific neutralization antibody.

**IMPORTANCE:** Exosomes participate in intercellular communnication between cells. Exosomes derived from virus-infected cells can mediate virus transmission or/and regulate immune response. However, the function of exosomes that from SVV-infected host cells during SVV transmission is unclear. Here, we demonstrate SVV can utilize host exosomes to establish productive infection in intercellular transmission. Furthermore, exosome-mediated SVV transmission is resistant to SVVV-specific neutralizing antibodies. This discovery sheds light on neutralizing antibodies resistant to SVVV transmission by exosomes as a potential immune evasion mechanism.

## INTRODUCTION

The Seneca Valley virus (SVV) is a single-stranded positive-strand RNA virus that belongs to the genus *Senecavirus* and family *Picornaviridae*. SVV has a typical icosahedral symmetry and a genome of 7.2 kb (1). In 2002, SVV was first discovered in the PER.C6 cell line in Maryland, USA (2). SVV mainly infects pigs, newborn piglets, fattening pigs, and other pigs of all ages, but neutralizing antibodies are also found in other animal bodies, such as cattle and sheep (3, 4). The clinical symptoms after SVV infection are very similar to the clinical symptoms of FMD. The main clinical symptoms are blisters and ulceration in the hoof and nose, as well as fever and anorexia (5). In recent years, the newly discovered SVV-induced blisters in pigs have brought great harm to China’s pig industry (6).

Exosomes are small vesicles with a diameter of 40–150 nm (7). Most of the model cells can secrete exosomes, which contain multiple substances, including large amounts of proteins and nucleic acids, and carry substances to a variety of cells (8-10). The virus enters the cell through endocytic pathways during the process of exosome formation and completes its own assembly and release (11). It has been reported in the literature that hepatitis A virus (a picornavirus) and hepatitis C virus can use exosomes to spread their DNA and escape the body’s immune response (12). However, the pathogenesis of SVV is not yet fully understood. In the view of the aforementioned background research, we suspect that exosomes may be an important mediator of SVV transmission between cells. Therefore, we aim to explore whether exosomes carry SVV for transmission and if the inhibition of exosome secretion and production inhibits the proliferation of SVV in cells.

In the present study, we extracted exosomes from SVV-infected IBRS-2 cells and identified them. We then introduced the extracted exosomes into 293T and IBRS-2 cells and found that SVV carried by exosomes can proliferate in these cells. We also inhibited IBRS-2 secretion and production of exosomes, which resulted in the inhibition of SVV proliferation. Interestingly, we also found that SVV carried by exosomes cannot be neutralized by SVV neutralizing antibodies. This study can provide an important reference for the pathogenesis of SVV and the antiviral mechanisms of the body.

## RESULTS

### Identification of exosomes extracted from SVV-infected IBRS-2 cells

Exosomes were extracted from SVV-infected IB-IR-2 cells, and the morphology of exosomes was identified by TEM. Further purification was performed with the use of a CD63-immunoaffinity kit. Cup-shaped lipid bilayer vesicles of representative exosome images were observed by TEM (Fig. 1a). To further identify exosomes by the exosome-associated protein markers, including Alix, CD63, and CD9, a WB experiment was performed. The exosomes extracted from SVV-infected IBRS-2 cells contained the exosome-associated protein markers Alix, CD9, and CD63 (Fig. 1b). At the same time, the size of exosomes was evaluated with the NTA method, and it was found that the particle size of the exosomes was mainly distributed in the range of 50150 nm (Fig. 1c). To verify whether the extracted exosomes contained SVV, we identified the SVV gene sequence in exosomes by qRT-PCR. SVV gene sequences were found in SVV-infected IBRS-2 cells and exosomes extracted from SVV-infected IBRS-2 cells (Fig. 1d). These results indicate that the morphology and particle size of the exosomes extracted in this study are consistent with those reported in the literature, and exosome-associated proteins are detected in exosomes. It is important that SVV is included in the exosomes.

**Fig 1:**
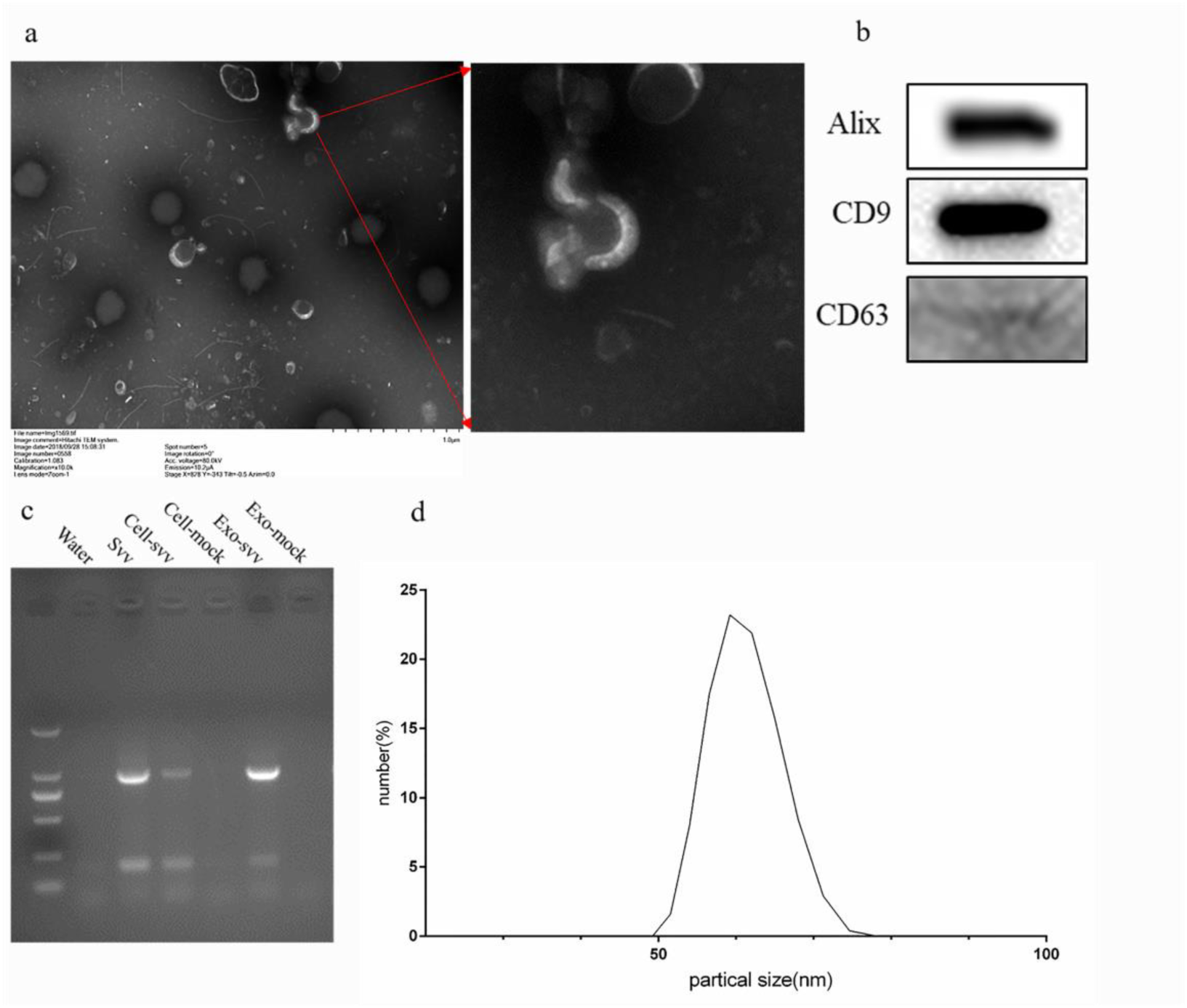
Isolation and characterization of exosomes extracted from SVV-infected IBRS-2 cells. (a) Transmission electron microscopic observation of exosomes extracted from SVV-infected IBRS-2 cells after negative staining with phosphotungstic acid. (b) Exosomes were extracted from SVV-infected IBRS-2 cells, purified, and identified by western blot with antibodies directed against Alix, CD9, and CD63. (c) SVV RNA genomic RNAs were identified in exosomes that were extracted from SVV-infected IBRS-2 cells (Exo-svv). At the same time, SVV, SVV-infected IBRS-2 cells (Cell-svv), normal IBRS-2 cells (Cell-mock), and exosomes extracted from normal IBRS-2 cells (Exo-mock) were used as controls. (d) Histogram displaying the mean size and size distribution profile of exosome particles isolated from the culture supernatants of SVV-infected IBRS-2 cells by the NTA method.

### Exosomes mediate the spread of SVV in susceptible and non-susceptible cells

In the aforementioned experiments, we found that the extracted exosomes contained SVV. It has been reported in the literature that exosomes can mediate the spread of substances in cells. Therefore, this study aimed to further explore whether exosomes mediate the spread of SVV. The DIL-stained exosomes were introduced into IBRS-2 and 293T cells, and at the same time, mock-exosomes(mock-exo) and SVV-GFP were introduced as the control. The fluorescence was observed under a laser confocal microscope. According to the experimental results, red fluorescence (DIL) was observed in both IBRS-2 and 293T cell membranes, and the GFP carried by SVV co-localized with DIL. In addition, only red fluorescence was observed in the mock-exo group (Fig. 2a). To further investigate whether exosomes mediate the spread of SVV, SVV-exosomes `(SVV-exo)` was added to IBRS-2 and 293T cells, and mock-exo and SVV-GFP were added as the control group. Finally, the copy number of SVV was detected by qRT-PCR. According to the experimental results, the copy number of SVV in the SVV-exo group was increased in both 293T and IBRS-2 cells compared with that in the mock group, and the copy number in IBRS-2 cells was higher than that in 293T cells (Fig. 2b). These results demonstrate that exosomes carry SVV to 293T and IBRS-2 cells, and SVV can proliferate in recipient cells.

**Fig 2:**
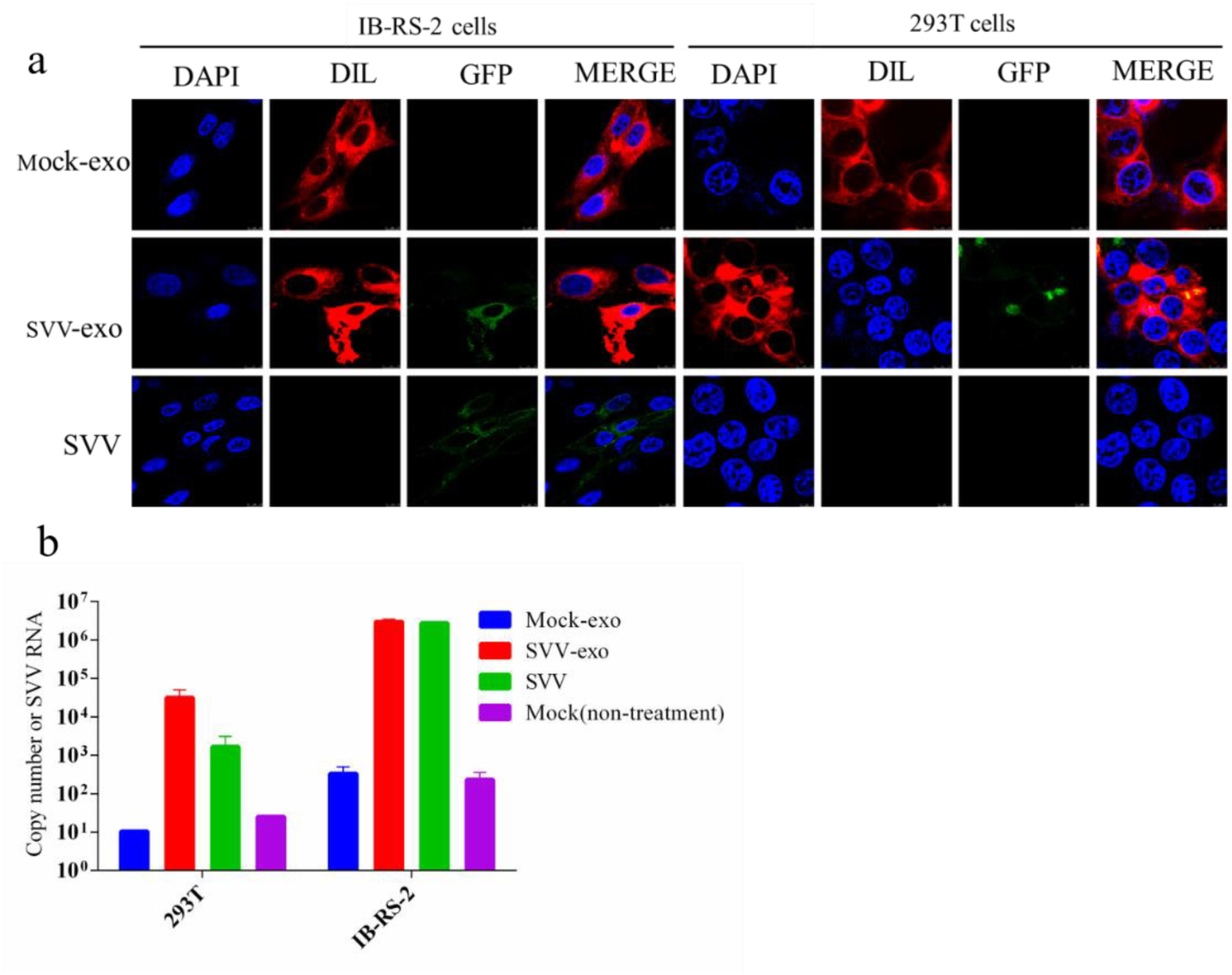
Exosomes mediate the spread of SVV in susceptible and non-susceptible cells. (a) Exosomes extracted from SVV-GFP-infected cells were stained with DIL, washed twice by ultracentrifugation, and incubated in 293T and IBRS-2 cells. Eight hours after incubation, the nuclei were stained with DAPI, and fluorescence was observed. (b) Exosomes extracted from SVV-infected IBRS-2 cells (SVV-exo) were incubated with 293T and IBRS-2 cells, and at the same time, exosomes extracted from normal IBRS-2 cells, SVV venom with the same viral load as SVV-exo, and PBS were used as controls. The virus copy number was detected by qRT-PCR 24 h after incubation.

### Inhibition of exosome secretion can inhibit the proliferation of SVV in IBRS-2 cells

In aforementioned experiments, we found that exosomes carried SVV and that SVV proliferated in 293T and IBRS-2 cells. Therefore, we next investigated whether inhibiting the secretion of exosomes affected the replication of SVV. To verify whether intracellular RAB27a expression was downregulated after si-RAN27a transfection, we transfected RBA27a into IB cells and detected RAB27a miRNA levels by qRT-PCR. The results showed that the miRNA expression of intracellular RAB27a was significantly downregulated in si-RBA27a-transfected cells compared with that in the control group (Fig. 3a). Then, in order to investigate whether the number of exosomes in the transfected cells was changed, we examined the miRNA expression level of the exosome protein marker ALIX. The results showed that the expression of ALIX was significantly upregulated in RAB27a-transfected IBRS-2 cells compared with that in the control group (Fig. 3b). In addition, the miRNA expression of ALIX was significantly upregulated in si-RAB27a-transfected cells. Therefore, we next explored whether the observed changes in the number of exosomes affected the copy number of SVV in cells. qRT-PCR was performed to detect the copy number of intracellular and extracellular SVV in si-RAB27a-transfected cells infected with SVV. The results showed that the intracellular SVV copy number was significantly upregulated (Fig. 3c) and the extracellular SVV copy number was significantly downregulated (Fig. 3d) compared with those in the control group. To re-validate the secretion of extracellular exosomes after the inhibition of extracellular secretion, we measured the number of extracellular exosomes in si-RAB27a-transfected cells by NTA. The results showed that after RAB27a transfection, the number of exosomes decreased significantly relative to the control group (Fig. 4). The aforementioned results indicate that the inhibition of RAB27a expression leads to a significant increase in the number of exosomes, and the copy number of intracellular SVV is also significantly upregulated.

**Fig 3:**
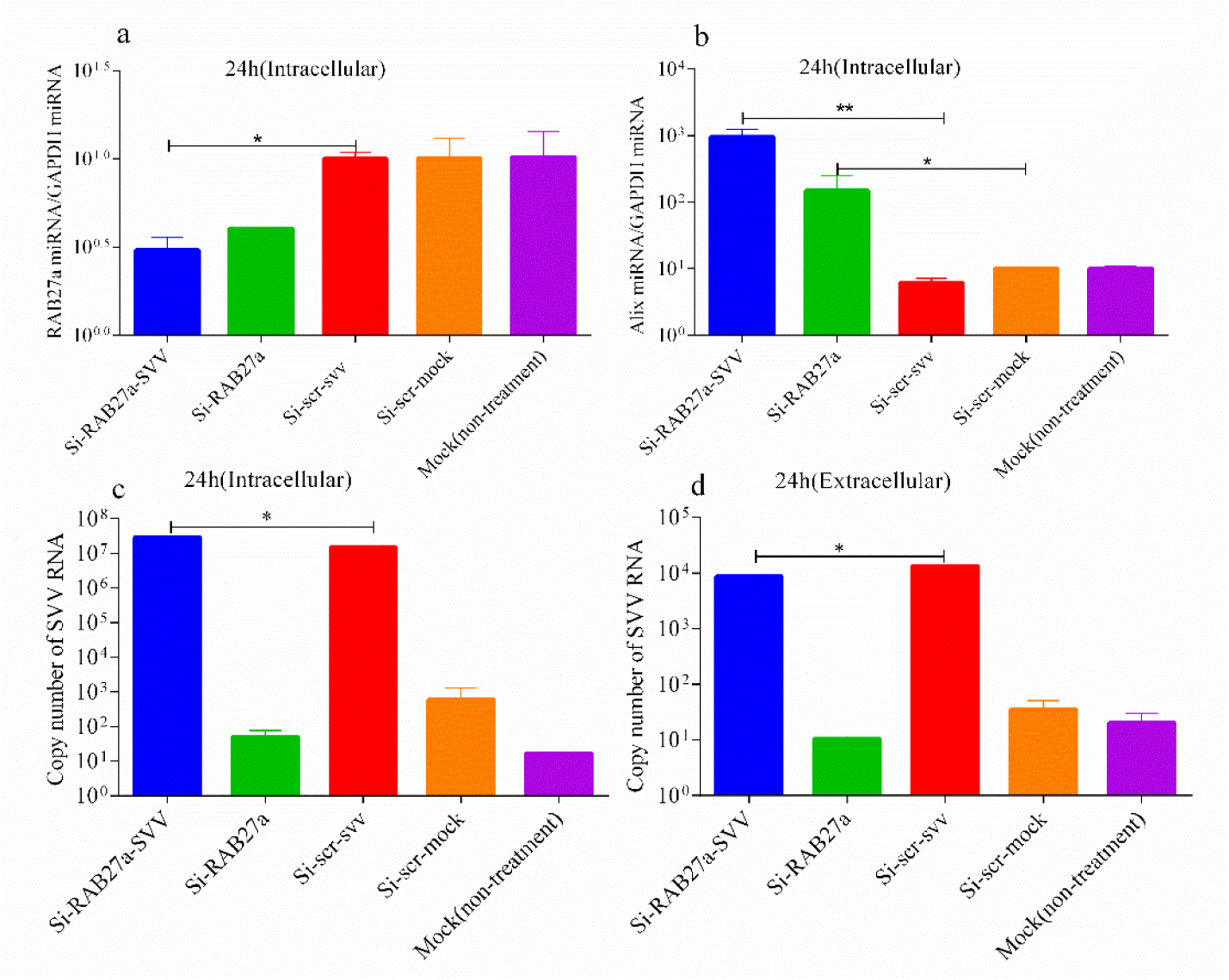
Changes in copy number of SVV. (a) Si-RAB27a was transfected into IBRS-2 cells, which were also infected with SVV, and the expression of RAB27a mRNA in IBRS-2 cells was detected by qRT-PCR. (a) Si-RAB27a was transfected into IBRS-2 cells, which were also infected with SVV, and the expression of Alix mRNA was detected by qRT-PCR. (c) Si-RAB27a was transfected into IBRS-2 cells, which were also infected with SVV, and the copy number of intracellular SVV in IBRS-2 cells was detected by qRT-PCR. (c) Si-RAB27a was transfected into IBRS-2 cells, which were also infected with SVV, and the copy number of extracellular SVV in IBRS-2 cells was detected by qRT-PCR. Significance was calculated using a two-tailed t-test and labeled as \**P* < 0.05 and \*\**P* < 0.01 in graphs.

**Fig 4:**
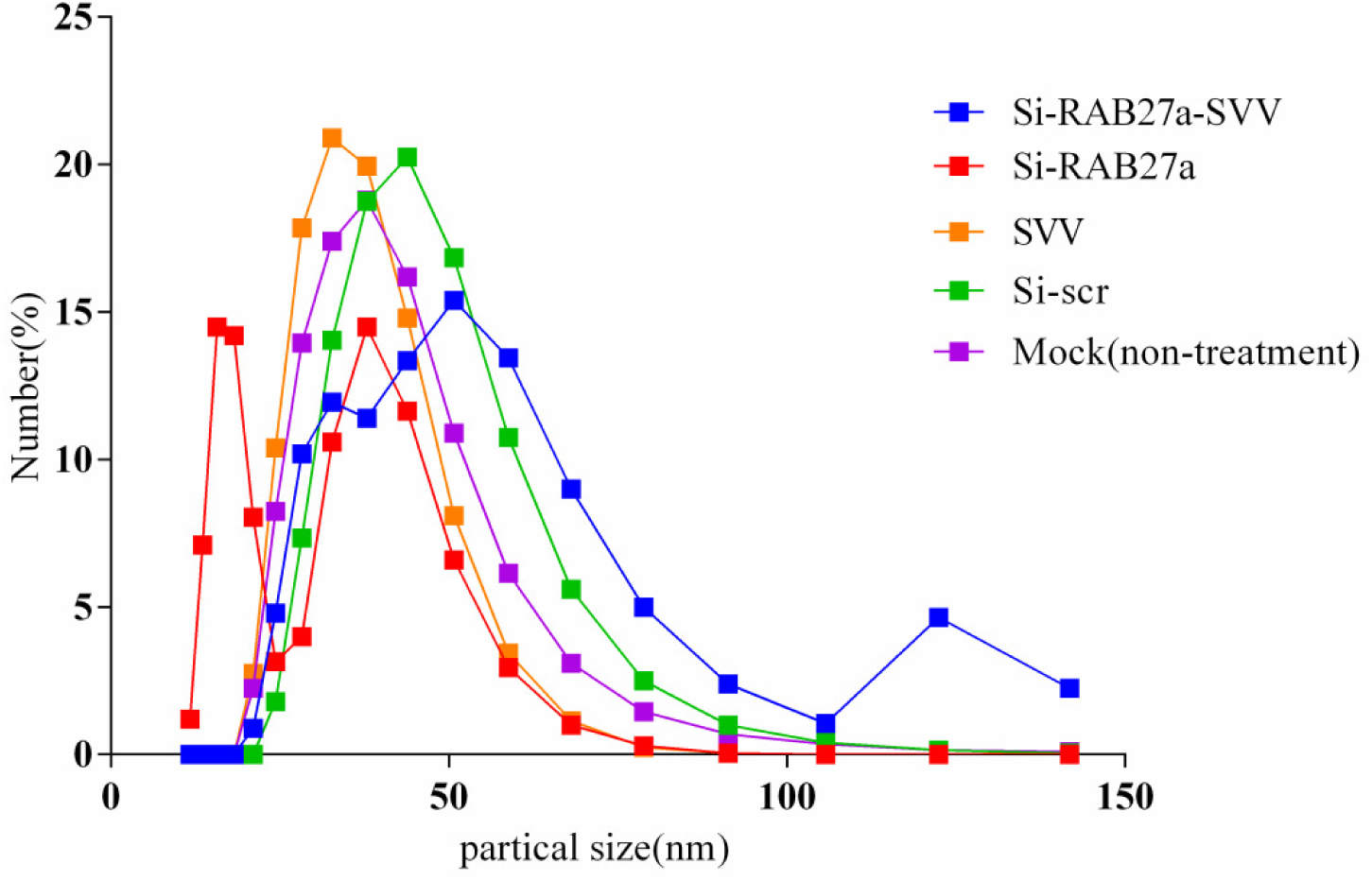
The number of exosomes secreted by IBRS-2 cells was detected after interfering with RAB27a. Si-RAB27a (100 pmol) was transfected into IBRS-2 cells, which were also infected with SVV. Exosomes were extracted from the culture supernatants of SVV-infected IBRS-2 cells, and the number of exosomes was detected by the NTA method.

### Inhibition of SVV proliferation by inhibiting the production of exosomes

In order to further verify the aforementioned experimental results, we determined whether the inhibition of exosome production inhibits the proliferation of SVV. IB-IS-2 cells were infected with SVV, the infected cells were treated with exosomes secretion inhibitor GW4869, and the copy number of intracellular and extracellular SVV was detected by qRT-PCR. According to the experimental results, the copy number of intracellular and extracellular SVV was significantly decreased after GW4869 treatment compared with that of the control group. The aforementioned results indicate that the inhibition of exosome production can inhibit the replication of SVV (Fig. 5).

**Fig 5:**
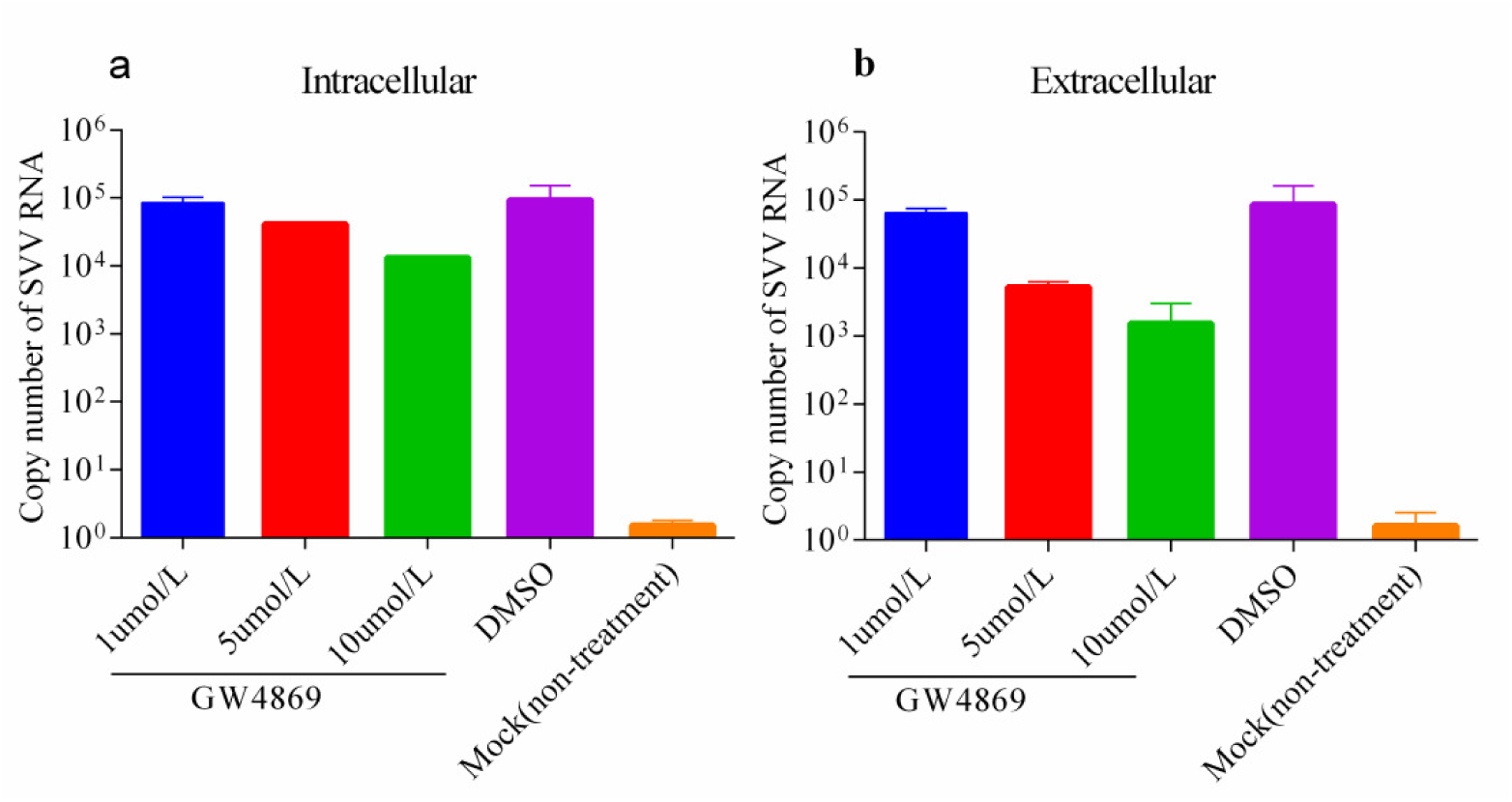
Inhibition of exosome release impairs SVV transmission mediated by exosomes. IBRS-2 cells were infected with SVV for 1.5 h and then incubated with 1, 5, and 10 µmol of GW4869 for 36 h. DMSO was used as a control, and cells without any treatment were used as a negative control. (a) The intracellular SVV copy number was detected. (b) The extracellular SVV copy number was detected.

### Exosomes promote the proliferation of SVV in susceptible cells

To further investigate whether exosomes extracted from IBRS-2 cells can promote SVV proliferation, we measured the proliferation of SVV in IBRS-2 cells with exosomes by qRT-PCR. The results showed that the SVV copy number of the mock-exo and SVV-exo groups was significantly higher than that of the SVV control group at 24 h after SVV infection (Fig. 6a), whereas 48 h after SVV infection, the copy number was higher only in the SVV-exo group (Fig. 6b). When the exocytosis dose was 50 ng, the SVV copy number of the mock-exo and SVV-exo groups was higher than that of the SVV control group at 24 and 48 h after SVV infection, but the number of SVV copies in the SVV-exo group was only significantly higher than that in the SVV control group 48 h after SVV infection (Fig. 6c and 6d).

**Fig 6:**
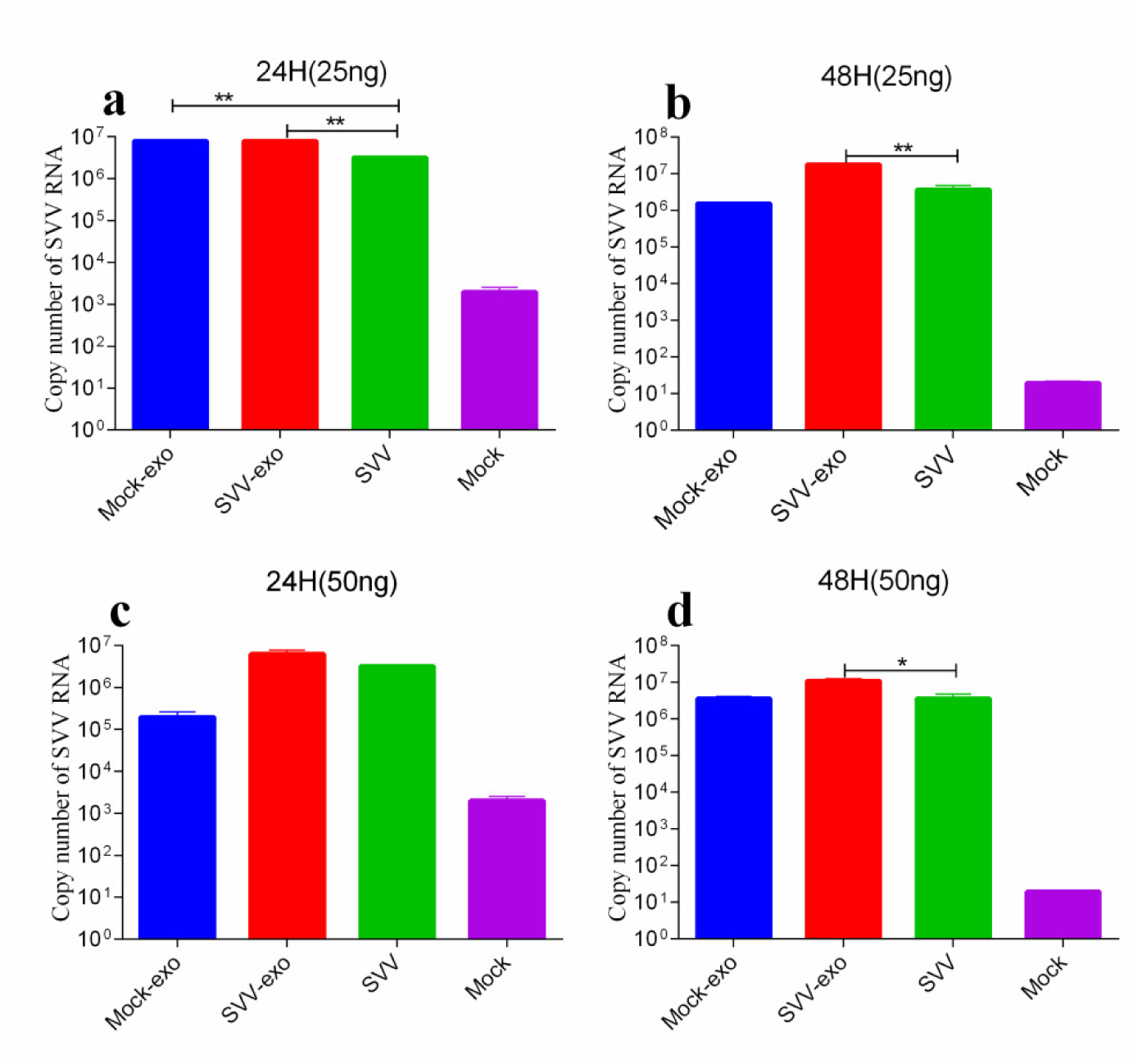
Exosomes promote the proliferation of SVV in IBRS-2. Exosomes were extracted from SVV-infected IB cells and cells without any treatment, and the protein concentration was measured. The extracted exosomes were divided into a high-dose group and a low-dose group and then re-inoculated into IB-SI-2 cells. After 8 h of incubation, exosomes were replaced with SVV, cells and culture supernatants were collected at 24 and 48 h after inoculation, and the copy number of SVV was detected. (a) The exosome protein concentration was inoculated at a dose of 25 ng, and the viral load was measured 24 h after SVV infection. (b) The exosome protein concentration was inoculated at a dose of 25 ng, and the viral load was measured 48 h after SVV infection. (c) The exosome protein concentration was inoculated at a dose of 50 ng, and the viral load was measured 24 h after SVV infection. (d) The exosome protein concentration was inoculated at a dose of 25 ng, and the viral load was measured 48 h after SVV infection. Significance was calculated using a two-tailed t-test and labeled as \**P* < 0.05 and \*\**P* < 0.01 in graphs.

### Exosome-mediated SVV infection is not inhibited by SVV NAbs

Because SVV is carried in exosomes, we investigated if the SVV contained in exosomes can be neutralized by SVV neutralizing antibodies. Exosomes extracted from SVV-infected IBRS-2 cells and SVV were diluted 10 times and then incubated with an SVV neutralizing antibody for 1.5 h. Exosomes and SVV were inoculated into IBRS-2 cells, and the copy number of SVV in the inoculated cells was detected by qRT-PCR (Fig. 7).

**Fig 7:**
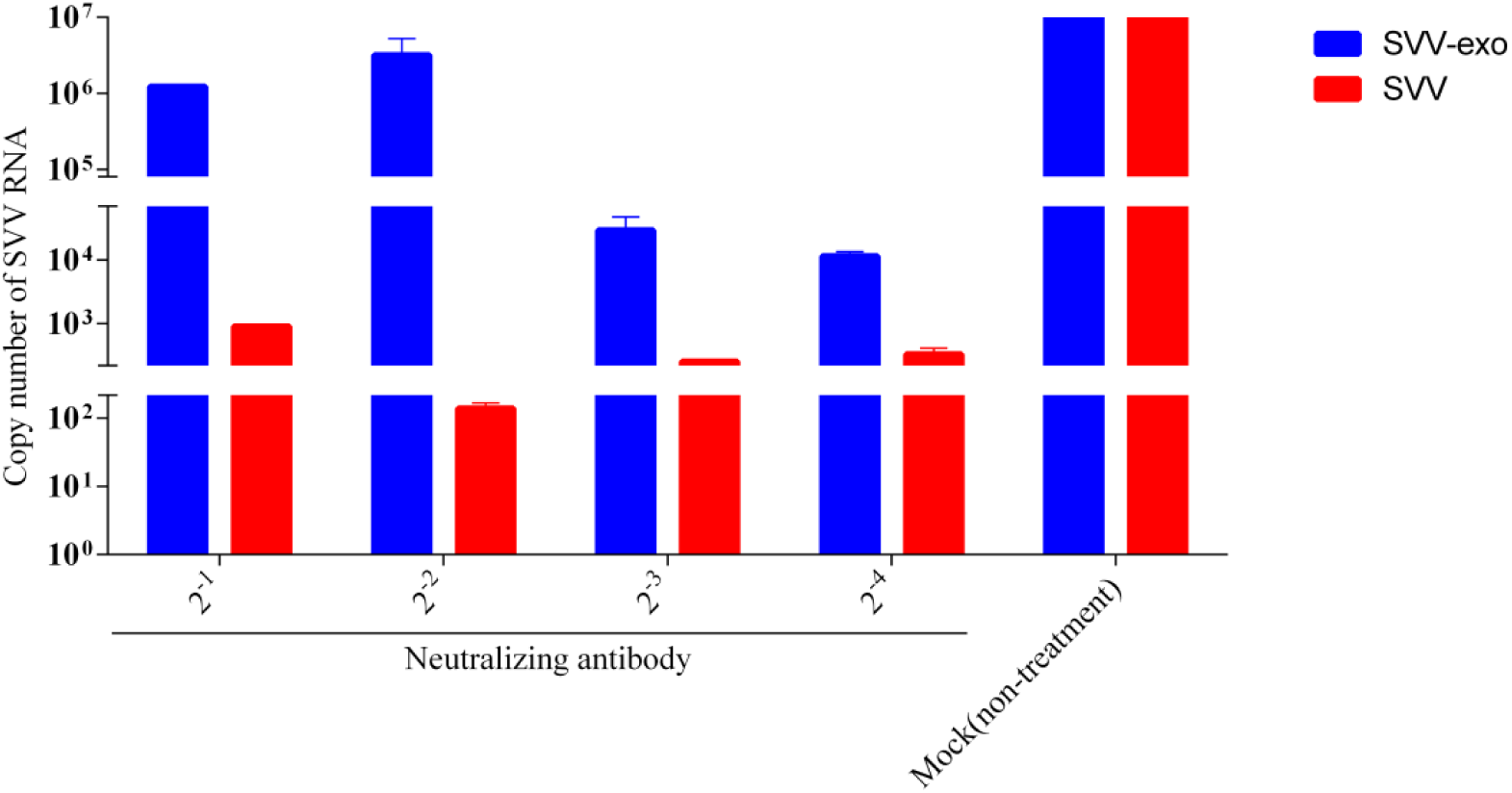
Exosome-mediated SVV infection is not blocked by SVV-specific neutralizing antibodies (NAbs). Purified SVV-positive exosomes and SVV were diluted and then the diluted exosomes and SVV were incubated with SVV-specific NAbs separately (the titer of serum neutralization against SVV was >1:1024, determined by VNT) for 1 h. Exosomes and SVV without any treatment were used as a control. Then, PK-15 cells were exposed to the NAbs-treated exosomes or SVV for 2 h. The exosomes or viruses were washed off with PBS at 37 °C, and the medium was replaced with fresh maintenance medium for 24 h. qRT-PCR was used to evaluateSVV replication in IBRS-2 cells.

## DISCUSSION

Exosomes are small vesicles that are secreted by cells (13). At present, there is no fixed standard for the separation and identification of exosomes. Many different methods are used for the isolation and identification of exosomes (14), such as UC, gradient UC, co-precipitation, size-exclusion chromatography, NTA electron microscopy (EM) analysis, and protein identification by CD63, CD9, CD81, and ALIX expression. However, UC combined with exosome marker protein-labeled magnetic bead purification is considered the gold standard method for exogenous separation and purification (7, 14, 15).

In this study, exosomes were extracted from SVV-infected and normal IBRS-2 cells, and the exosomes were identified by WB, NTA, TEM, and qPCR. According to the results, the extracted exosomes contained CD63, CD9, and ALIX, which are protein markers of exosomes. In addition, their diameters were about 40–150 nm, and the cup-like structure of the membrane was observed using TEM. The identification of exosomes extracted from the IBRS-2 cells was consistent with the results that are reported in the literature. Furthermore, the exosomes extracted from the cells of the SVV-infected group contained the SVV nucleic acid sequence, which was determined by qPCR.

In the current study, we suggest that exosomes can serve as a delivery vector for pathogen-associated molecules, which help pathogenic microorganisms to spread infections throughout the body’s microenvironment (16). In addition, their role in viral infections is receiving more and more attention (17, 18). Some viruses are included in the exosomes for transmission, which may be an important way to escape the immune response (19). During the process of exosome formation, some viruses enter the cell through exosomes, mainly in the endocytic pathway, which can deliver the virus directly into the cell without the need for cell membrane receptors (12, 19, 20). In our study, we isolated exosomes with DIL staining and introduced them into IBRS-2 and 293T cells. We found that the exosomes were able to re-enter the cells, validating the activity of the exosomes. Then, we inoculated SVV and exosomes with the same copy number of SVV into IBRS-2 and 293T cells and found that SVV contained within exosomes proliferated in IBRS-2 and 293T cells. However, the copy number in 293T cells was lower than that in IBRS-2 cells. Therefore, exosomes extracted from IB cells can enter IB and 293T cells and mediate the spread of SVV.

The aforementioned results suggest that exosomes can mediate the spread of SVV, but do exosomes play an important role in the spread of SVV in `host` cells? Furthermore, does the inhibition of exosome secretion affect the proliferation of exosomes? Studies have shown that silencing the expression of Rab27a and Rab27b reduces exosome secretion of CD63, CD81, and MHC class II, and the downregulation of Rab27 effector Slp4 and Slac2b also inhibits exosome secretion (21, 22). We synthesized RAB27a siRNA and transfected it into IBRS-2 cells and then detected the expression level of RAB27a miRNA and ALIX. The number of intracellular exosomes was significantly upregulated after the downregulation of RAB27a expression. The copy number of intracellular SVV increased significantly, whereas the copy number of extracellular SVV decreased significantly after RAB27a interference. GW4869 inhibits exosome formation by inhibiting neutral sphingomyelinase 2 (23). We treated SVV-infected IBRS-2 cells with different doses of GW4869 and measured the copy number of SVV *in vitro* and *in vivo*. Unlike with si-RAB27a, the number of copies of intracellular SVV decreased in a dose-dependent manner after the treatment of cells with GW4869. According to our current experimental results, it cannot be explained why the intracellular SVV copy number was significantly increased after RAB27a interference but decreased after GW4869 treatment. We hypothesize that this may be related to the different mechanisms by which these two inhibitors block exosome formation and secretion. In any case, the copy number of extracellular SVV was decreased following the treatment with these two different methods.

*In vitro* infected EV71-isolated exosomes are rich in miRNA-146a, and miRNA-146a can inhibit type I interferon and promote EV71 replication (24). In our study, exosomes were found to promote SVV replication. We treated IB-RS cells with SVV-exosomes and mock-exo at the same protein dose and then inoculated SVV. We found that after treatment with SVV-exosomes and mock-exosomes at 24 and 48 h of SVV infection, the copy number of SVV was higher than that of SVV infection, and it was more significant at 24 h. Our current research is insufficient to explain how exosomes promote the proliferation of SVV.

Studies have shown that viruses carried by exosomes, such as PRRSV, HCV, and EV71, cannot be neutralized by neutralizing antibodies (25-30). It is well established that some infected pigs have high levels of antibodies, but the virus in the body can be detected, which may be related to the protection of exosomes. According to our current research, SVV carried in exosomes was not neutralized by SVV neutralizing antibodies, which is consistent with the results reported in related literature. This also suggests an important way for the virus to enter into exosomes, thereby evading the body’s immune response.

In conclusion, we successfully extracted, purified, and identified exosomes from SVV-infected IBRS-2 cells and determined that exosomes can carry SVV, which allows the proliferation of the virus in susceptible and non-susceptible cells. The inhibition of exosome secretion and production inhibits the replication of SVV. Moreover, exosomes extracted from IBRS-2 cells can promote the proliferation of SVV. SVV carried in exosomes cannot be neutralized by SVV neutralizing antibodies. Taken together, these datas revealed an advanced and novel mechanism for better understanding that viral transmission through exosomes contributes to the known immune evasive properties of SVV.

## MATERIALS AND METHODS

### Cell culture

To obtain a cell culture supernatant for exosome extraction, we used IBRS-2 cells as a model. IBRS-2 cells were cultured in Dulbecco’s modified Eagle’s medium (DMEM) supplemented with 10% fetal bovine serum (FBS), 100 IU/mL penicillin, and 100 mg/mL streptomycin. The cells were cultured in an incubator maintained at 37 °C and with a CO_2_ concentration of 5%.

### SVV infection and cell culture supernatant collection

In order to obtain exosomes secreted by SVV-infected cells, we inoculated SVV into IBRS-2 cells and collected the supernatants at specific times after infection. SVV was isolated previously, as described later in the text, and preserved by our lab (China Reference Laboratory Network for FMD) (31). IBRS-2 cells were incubated in a 150-mm culture dish until they became confluent (Corning, New York, USA). The culture supernatant was then discarded, the cells were washed with PBS, and FBS-free DMEM was added. SVV (0.05 TCID50) was inoculated, and PBS was used as a control. After 1 h of incubation, SVV was discarded and replaced with DMEM containing 2% exosome-depleted FBS. The cell culture supernatant was collected after 36 h of culture.

### Exosome isolation and purification

In order to further separate and purify the collected supernatant, we performed differential centrifugation with the collected supernatant. All of the following centrifugation processes were carried out in a 4 °C environment. The collected supernatant was centrifuged at 500 × *g* for 5 min to remove larger fragments and cells, and then the supernatant was collected and centrifuged at 2,000 × *g* for 10 min to further remove cell debris. To remove any cells, the collected supernatant was centrifuged at 12,000 × *g* for 45 min. The large vesicles were collected and filtered through a 0.22-µm filter. Finally, the collected supernatant was centrifuged at 120,000 × *g* for 2 h with an ultracentrifuge (Thermo Scientific Sorvall WX100), and the contents at the bottom of the centrifuge tube were resuspended in 500 μL of PBS. To further purify the extracted exosomes, we used a CD63 antibody-labeled exosomes isolation kit (Miltenyi Biotec, Bergisch Gladbach, Germany).

### Transmission electron microscopy (TEM)

Direct morphological observation of the characteristics of exosomes is an important method for exosome identification (Shao et al., 2018). Therefore, we analyzed the extracted exosomes using TEM (Hitachi H-7000FA, Tokyo, Japan). After observation, we first extracted the exosomes with a TEM 200 copper mesh (EMS 80100-Cu US) and then stained the exosomes with phosphoric acid dock for 2 min. After drying under an incandescent lamp, the electron microscope was used to observe the extracted exosomes, and the observed voltage was 80 kV.

### Western blot analysis

Western blot (WB) analysis was performed using the following protocols. Briefly, purified exosomes were lysed with a radio-immunoprecipitation assay buffer (Santa Cruz Biotechnology, Dallas, TX, USA), and the cleared lysate was collected by centrifugation for protein separation on 12% sodium dodecyl sulfate-polyacrylamide gel electrophoresis gels. After electrophoresis, the separated proteins were transferred onto 0.45 μm polyvinylidene difluoride membranes (Millipore, USA). Next, the membranes were blocked for 1 h with 10% fat-free milk in Tris-buffered saline containing Tween 20 (TBST). The blots were then incubated with primary antibodies at 4 °C overnight. Primary antibodies for CD63 (Abcam, Cambridge, UK), CD9 (Abcam, Cambridge, UK), and Alix (Cell Signaling Technology, Waltham, MA, USA) were used. After washing three times with TBST, the membranes were incubated with horseradish peroxidase-labeled secondary antibodies (Proteintech, Chicago, IL, USA) for 2 h at room temperature. Finally, the proteins were visualized with a clarity enhanced chemiluminescence WB substrate (Bio-Rad Laboratories, Hercules, CA, USA).

### Analysis and quantification of SVV RNA

For the PCR detection of SVV RNA, total RNA from exosomes and cells was extracted with a total exosome RNA and protein isolation kit (Life Technologies, USA) according to the manufacturer’s instructions. To quantify the RNA copies of SVV in SVV-infected or exosome-treated cells, total RNAs from cell culture samples were isolated with the E.Z.N.A. total RNA kit I (Omega Bio-tek). Detection of the number of copies of extracted RNA was performed using the Real-Time One-Step RT-PCR reagent (Takara). The following was the reaction system: 2X One-Step RT-PCR Buffer III 10 μL, TaKaRa Ex Taq HS (5 U/μL) 0.4 μL, Prime Script RT Enzyme Mix II μL, PCR forward primer (10 μM) 0.4 μL, PCR reverse primer (10 μM) 0.4 μL, SVV-3D probe 0.8 μL, total RNA 2 μL, and RNase-free dH_2_O 5.2 μL (PCR primers and the SVV-3D probe were provided by our laboratory). The reaction times and temperatures of the PCR were 42 °C for 15 min (1 cycle) and 40 cycles of 94 °C for 10 s, 57 °C for 30 s, and 72 °C for 30 s. The Applied Biosystems 7300 Real-Time PCR System (Thermo Fisher) was used.

### Nanoparticle tracking analysis (NTA)

The mean size and size distribution profile of exosomes that were isolated and purified from SVV-infected or control-treated IBRS-2 cell culture supernatants were analyzed, as described previously (26, 32). In brief, exosomes were diluted 100-fold with PBS prior to analysis, and the relative concentration was calculated on the basis of the dilution factor. Data analysis was performed with NTA 3.2 software (Malvern Panalytical Ltd., Malvern, Worcestershire, UK), and the samples evaluated using a Nanosight NS300 instrument (Malvern Panalytical Ltd., Malvern, Worcestershire, UK). Each sample was analyzed five times, and the counts were averaged.

### Fluorescence localization of exosomes

The exosomes extracted from SVV-green fluorescence protein (GFP)-infected cells and normal cells were purified using an exosome purification kit (Miltenyi Biotec, Bergisch Gladbach, Germany), and the copy number of SVV was detected. The exosomes were stained with DIL and washed twice by ultracentrifugation (UC). IBRS-2 and 293T cells were plated in a 20 mm laser confocal dedicated cell culture dish (Thermo Scientific Nunc). Exosomes and SVV-GFP with the same number of SVV copy numbers were inoculated into cells and incubated for 8 h. The cells were then fixed with paraformaldehyde, stained with DAPI, and sealed to avoid light during the aforementioned experimental process. The uptake of the fluorescently labeled SVV in IBRS-2 and 293T cells was visualized with a fluorescence microscope (Leica, Germany).

### Si-RAB27a transfection and quantification of RAB27a and Alix miRNA

RAB27a interfering RNA (100 pmol) was transfected into IBRS-2 cells using liposome 2000. SVV was inoculated 24 h after transfection, and cells were harvested 24 h after SVV inoculation. Total RNAs from cell culture samples were isolated with the E.Z.N.A. total RNA kit I (Omega Bio-tek). The reverse transcription of RNA into cDNA was performed using a Prime Script TM RT Master Mix (Takara). The following was the reaction system: 5X PrimeScript RT Master Mix (Perfect Real Time) 2 μL, total RNA 5 μL (200 ng), and RNase-free dH_2_O 3 μL. The reaction temperature and time of the PCR were 37 °C for 15 min and 85 °C for 5 s (reverse transcription reaction), respectively. Using GAPDH as an internal reference gene, qRT-PCR was performed using TB Green TM Premix Ex TaqTM II (Takara). The following was the reaction system: TB Green Premix Ex Taq II (Tli RNaseH Plus) 10 μL, PCR forward primer (10 μM) 0.8 μL, PCR reverse primer (10 μM) 0.8 μL, DNA (<100 ng) 2 μL, and sterilized water 6.4 μL.

### Exosome treatment with a SVV-specific neutralizing antibody

IBRS-2 cells were plated into 12-well cell culture plates, and the cells were replaced with serum-free DMEM once the cells reached 70%–80% confluency. Exosomes extracted from SVV-infected cells and SVV were simultaneously diluted to obtain concentrations ranging from 10^−1^ to 10^−4^. The diluted exosomes and SVV were incubated with the SVV neutralizing antibody for 1.5 h at 37 °C and then added to the prepared IBRS-2 cells. At the same time, exosomes and SVVs that were not incubated with the SVV neutralizing antibody were used as controls. The cells were cultured in a 5% CO_2_ cell culture incubator at 37 °C for 24 h. The cells and culture supernatants were used to detect the copy number of SVV.

## ACKNOWLEDGMENTS

This work was supported by grants from the Science and Technology Major Project of Gansu Province (19ZD2NA001).

